# Spatio-temporal requirements of Aurora kinase A in mouse oocytes meiotic spindle building

**DOI:** 10.1101/2024.04.01.587547

**Authors:** Cecilia S. Blengini, Michaela Vaskovicova, Jan Schier, David Drutovic, Karen Schindler

## Abstract

Meiotic spindles are critical to ensure proper chromosome segregation during gamete formation. Oocytes lack centrosomes and use alternative microtubule nucleation pathways for spindle building. However, how these mechanisms are regulated is still unknown. Aurora kinase A (AURKA) is necessary and sufficient for oocyte meiosis in mouse because *Aurka* KO oocytes arrest in meiosis I [1] and AURKA compensates for loss of *Aurkb*/*Aurkc* [2]. AURKA is required early in pro-metaphase I to trigger microtubule organizing center fragmentation, a step necessary to effectively build a bipolar spindle. Moreover, in double *Aurkb*/*Aurkc* knockouts, AURKA localizes to spindles and chromatin to support meiosis. Although these mouse models were useful for foundational studies, we were unable to resolve AURKA spatial and temporal functions. Here we provide high-resolution analyses of AURKA requirements during multiple steps of meiotic spindle building and identify the subcellular populations that carry out these functions. By combining mouse genetics and pharmacological approaches we show that AURKA is specifically required in early spindle building and later for spindle stability, whereas AURKC is specifically required in late pro-metaphase. Through expression of targeted AURKA constructs expressed in triple Aurora kinase knockout oocytes and high-resolution live imaging, we demonstrate that the spindle pole population of AURKA is the predominate pool that controls meiotic spindle building and stability.

## Results and discussion

Oocyte aneuploidy is the leading cause of early miscarriage. Spindles are critical for ensuring euploidy and spindle mechanics are frequently faulty in oocytes [3]. Oocytes rely on two mechanisms to form bipolar meiotic spindles: the Ran-GTP pathway [4-7] and acentriolar microtubule organizing centers (aMTOCs) [5, 8, 9]. The Ran-GTP pathway generates a gradient of spindle assembly factors from chromatin to control poleward polymerization of microtubules (MTs) [6, 10]. aMTOCs nucleate microtubules and form spindle poles [8]. During mouse oocyte meiotic maturation, aMTOCs undergo a series of structural changes. First, upon nuclear envelope breakdown (NEBD), aMTOCs fragment. Then, these fragments sort and separate on either side of chromosomes and cluster together to form two spindle poles [11]. How these processes are regulated is still not fully understood.

The Aurora protein kinases (AURK) have been extensively studied for the roles they play in regulating accurate chromosome segregation in mitosis and meiosis. The mammalian genome encodes three members: Aurora kinase A (AURKA), Aurora kinase B (AURKB), and Aurora kinase C (AURKC). AURKA mainly localizes with aMTOCs [12, 13], with a minor chromosomal population [14]. Through mouse knockout studies, we found that although each isoform is important for female fertility and oocyte meiosis, only AURKA is necessary and sufficient for fertility [1, 2]. Females that lack AURKB and AURKC (BC-knockout; KO), in which oocytes contain only AURKA, are fertile. In these oocytes, AURKA increases its population on chromosomes, compensating for loss of AURKB/C [2]. On the other hand, in oocyte-specific *Aurka* knockout mice, females are sterile, and their oocytes arrest at metaphase I with unfragmented aMTOCs and a disorganized TACC3-dependent spindle domain, a marker of a so-called liquid-like spindle domain (LISD). As a result, these oocytes have small, abnormal spindles [1]. aMTOC fragmentation is the first step of spindle assembly. It is unclear whether AURKA is required for later spindle formation steps, and which localized AURK population is essential for spindle formation. Here, we aimed to evaluate the spatial and temporal requirements of AURKA during meiotic maturation in mouse oocytes.

### AURKA is required for multiple steps of meiotic spindle formation

To temporally resolve specific AURKA requirements in oocyte meiosis, we inhibited AURKA with MLN8237 (MLN) in WT oocytes. The dose of MLN chosen was previously determined to be specific for AURKA [15]. We focused on determining AURKA functions in three critical spindle formation steps (Figure 1A). Each group was immunostained with Pericentrin (Pcnt), TACC3, and α-tubulin antibodies to visualize aMTOCs, LISD, and the spindle, respectively. We evaluated changes in aMTOC numbers and volume, TACC3 intensity, and spindle volume. When inhibiting middle and late steps, we examined oocytes prior to MLN-addition to ensure that the earlier processes had occurred normally. First, when we evaluated early pro-metaphase I (Figure 1B), MLN-treated oocytes had fewer aMTOCs, which were larger in volume than aMTOCs in control oocytes, consistent with fragmentation failure described in *Aurka* KO oocytes [1] (Figures 1C, D; S1A, B). Furthermore, spindle volume and TACC3 intensity measurements revealed two distinct clusters: MLN-treated oocytes had smaller spindles and reduced TACC3 intensity compared to controls (Figures 1C, E; S1C, D). These results are consistent with our previous data in *Aurka* KO oocytes [1].

**Figure 1.**
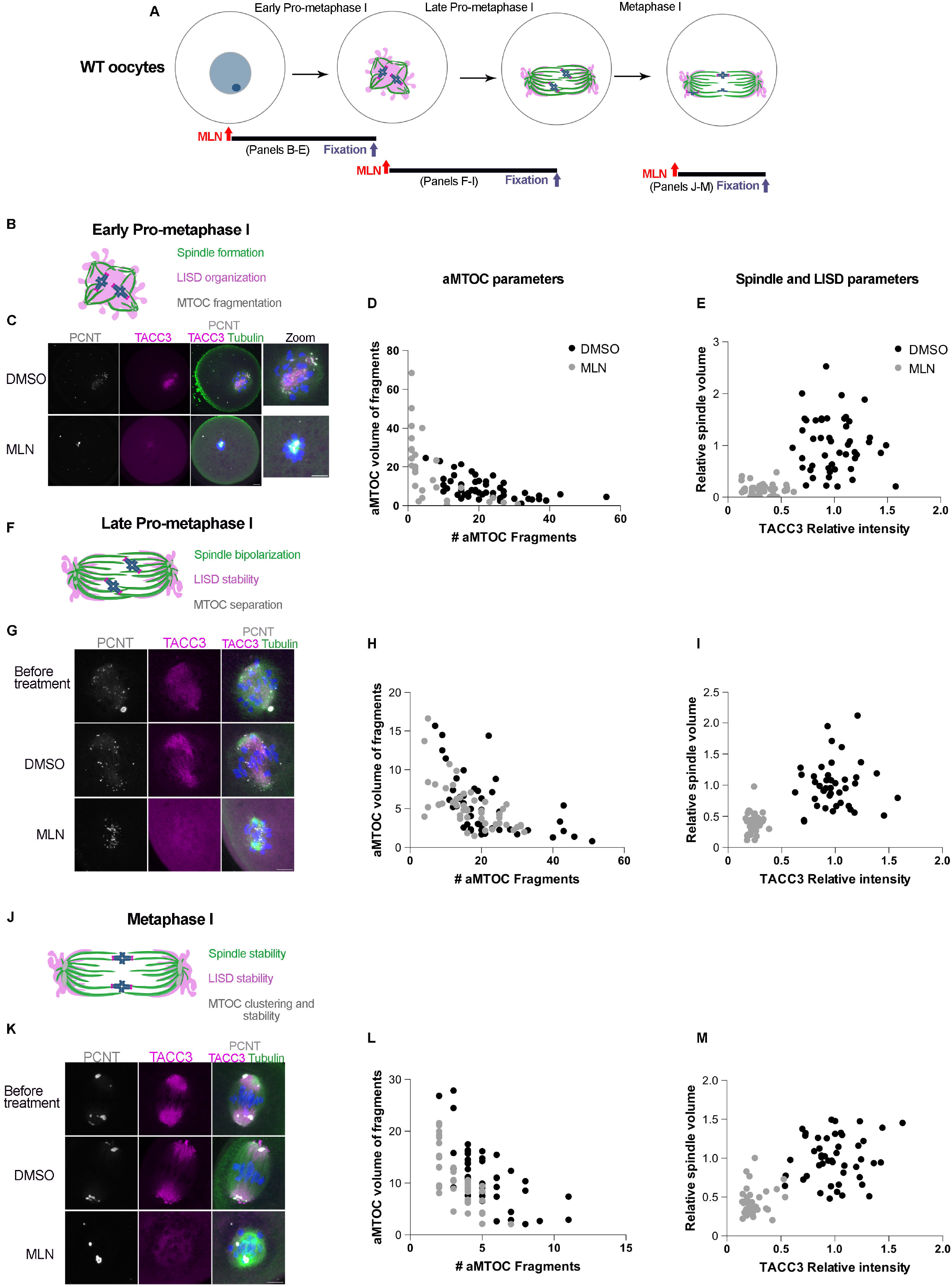
AURKA is required at early pro-metaphase I and metaphase I for spindle building. (A) Experimental design of WT oocytes treated with MLN during oocyte maturation. Red arrows indicate MLN addition time; blue arrows indicate fixation time. (B, F, J) Schematic of spindles at indicated stages, and the processes analyzed. (C, G, K) Representative confocal images of oocytes fixed at the indicated stages with and without MLN, immuno-stained with PCNT (gray), TACC3 (magenta), Tubulin (green) and DAPI (blue). Scale bars 10 µm. (D, H, L) Quantification of aMTOC parameters in (A), (G), (K) respectively. (E, I, M) Quantification of spindle and LISD parameters in (A), (G), (K) respectively. Black dots: DMSO-treated; gray dots: MLN-treated. See also Figure S1 which contains statistics.

In contrast, when AURKA was inhibited at late pro-metaphase I (Figure 1A, F), aMTOC number and volume were not altered compared to controls (Figures 1G, H; S1E, F). However, spindle volume and TACC3 intensity were different. Oocytes exposed to MLN produced smaller spindles (∼55% reduction) with reduced TACC3 intensity compared to controls (Figures 1G, I; S1G, H), suggesting that AURKA is essential to maintain LISD stability and spindle length.

Finally, we evaluated metaphase I spindle stability (Figure 1A, J) by inhibiting AURKA after bipolar spindles formed. Again, the data clustered into two groups. MLN-treated oocytes had fewer aMTOCs (Figure 1K, L; S1I, J) compared to controls, and they had a significant reduction in both spindle volume and TACC3 intensity (Figures 1K, M; S1K, L). Taken together, these data indicate that AURKA plays a critical role in several steps during the spindle formation process. First, AURKA controls aMTOC fragmentation and LISD organization, second, AURKA maintains LISD stability and spindle length, and third, AURKA is required for spindle stability by maintaining LISD organization and aMTOC clustering at poles.

AURKA and AURKC genetically interact to control spindle building [2], but which critical steps they function in are unclear. To fully understand how AURKA and AURKC participate in spindle building, we compared the differences of MLN treatment between WT (AURKA and AURKC) and BC-KO (AURKA only) oocytes (Figure 2A). When evaluating early pro-metaphase I and metaphase I spindles, we did not observe differences compared to WT oocytes treated with MLN (Figure 2B-E; J-M; S2A-D; I-L). However, when we examined late pro-metaphase I (Figure 2F), MLN-inhibited BC-KO oocytes behaved differently from WT. BC-KO oocytes treated with MLN had over-clustered aMTOCs, characterized by fewer fragments with greater volume than in controls (Figures 2G, H; S2E, F). Moreover, the TACC3 domains were also disorganized, and the spindle volume was reduced by 70%, compared to 55% in WT oocytes (Figures 2G, I; S2G, H). These differences between WT and BC-KO oocytes suggest a role for AURKC to separate aMTOCs during late-prometaphase I, a function which needs further investigation.

**Figure 2.**
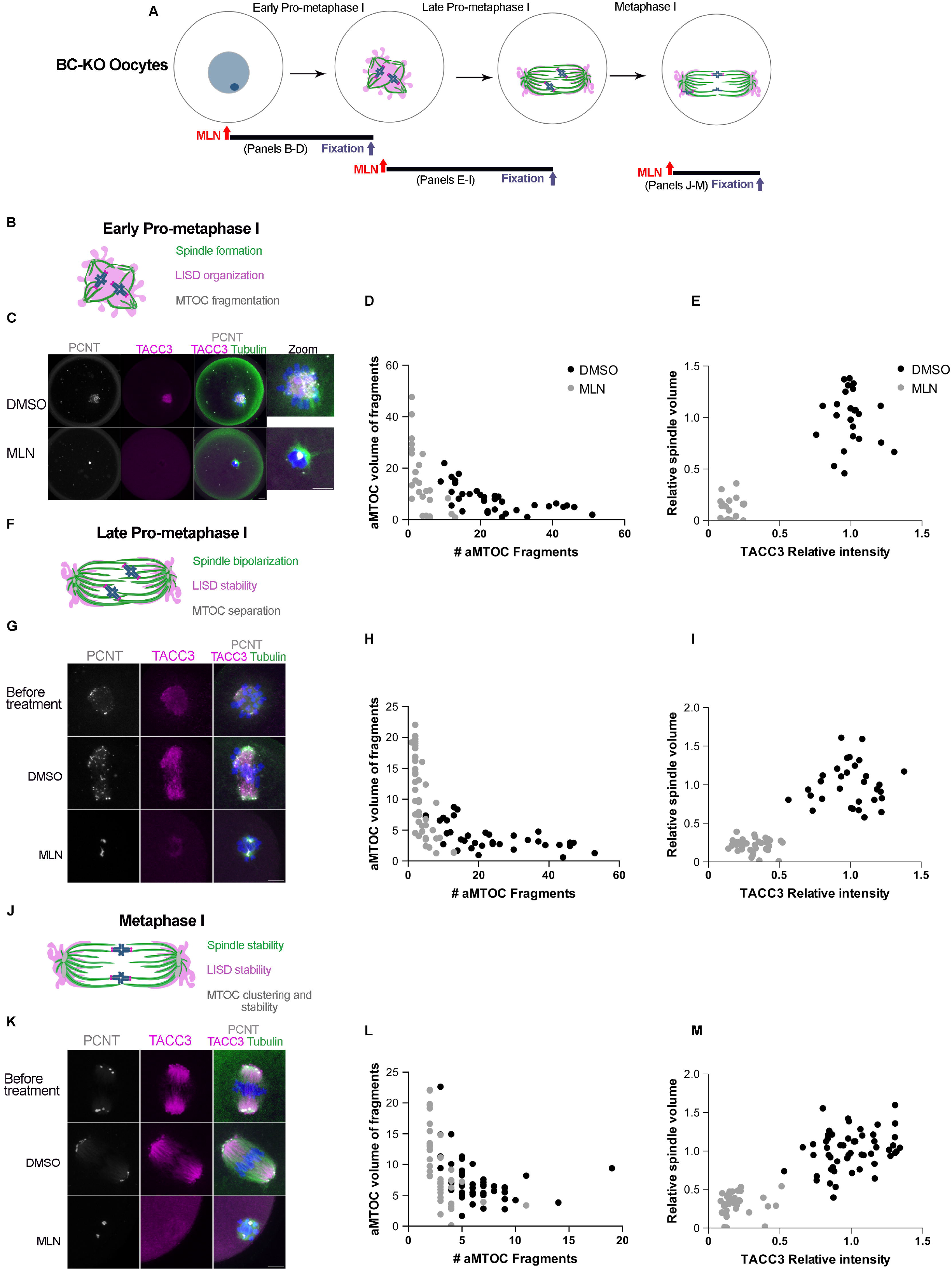
AURKC is required at late pro-metaphase I for aMTOC separation. (A) Experimental design of BC-KO oocytes treated with MLN; Red arrows indicate MLN addition time; blue arrows indicate fixation time. (B, F, J) Schematic of spindles at the indicated stages, and the processes analyzed. (C, G, K) Representative confocal images of oocytes fixed at the indicated stages with and without MLN, immuno-stained with PCNT (gray), TACC3 (magenta), Tubulin (green) and DAPI (blue). Scale bars 10 µm. (D, H, L) Quantification of aMTOC parameters in (A), (G), (K), respectively. (E, I, M) Quantification of spindle and LISD parameters in (A), (G), (K), respectively. Black dots: DMSO-treated; gray dots: MLN-treated. See also Figure S2 which contains statistics.

We observed different responses to AURKA inhibition between WT and BC-KOs. To refine the temporal order in spindle building steps, we acutely treated early pro-metaphase I oocytes with MLN and performed live-cell imaging using light sheet microscopy. Prior to adding MLN, we observed an increase in spindle volume and an increase aMTOC number in both WT and BC-KO oocytes (Video S1). Compared to controls, after MLN addition in both WT and BC-KO oocytes, spindle volume rapidly reduced (10 min) (Figure 3A-B, D). Notably, 56% of BC-KO oocytes lost their spindles, a phenotype not observed in WT MLN-treated oocytes (Figure 3A; Video S1). The behavior of aMTOCs was also altered. Both WT and BC-KO MLN-treated oocytes had fewer aMTOCs that were slightly larger (Figure 3A, C, E). In BC-KO oocytes aMTOC reduction was greater compared to WT oocytes. These results suggest that AURKA is specifically required in early spindle building and later for spindle stability, whereas AURKC is specifically required in late pro-metaphase to regulate aMTOCs. One possibility is that AURKC has a role in separating and clustering aMTOCs at late pro-metaphase I. Previous evidence showed that loss of AURKC activity in mouse oocytes causes defects in aMTOC clustering resulting in multipolar spindles [16]. However, BC-KO oocytes may be more susceptible to AURKA inhibition than WT, because the localized amounts of AURKA protein are now reduced to compensate in other subcellular localizations where AURKB/C once were [2].

**Figure 3.**
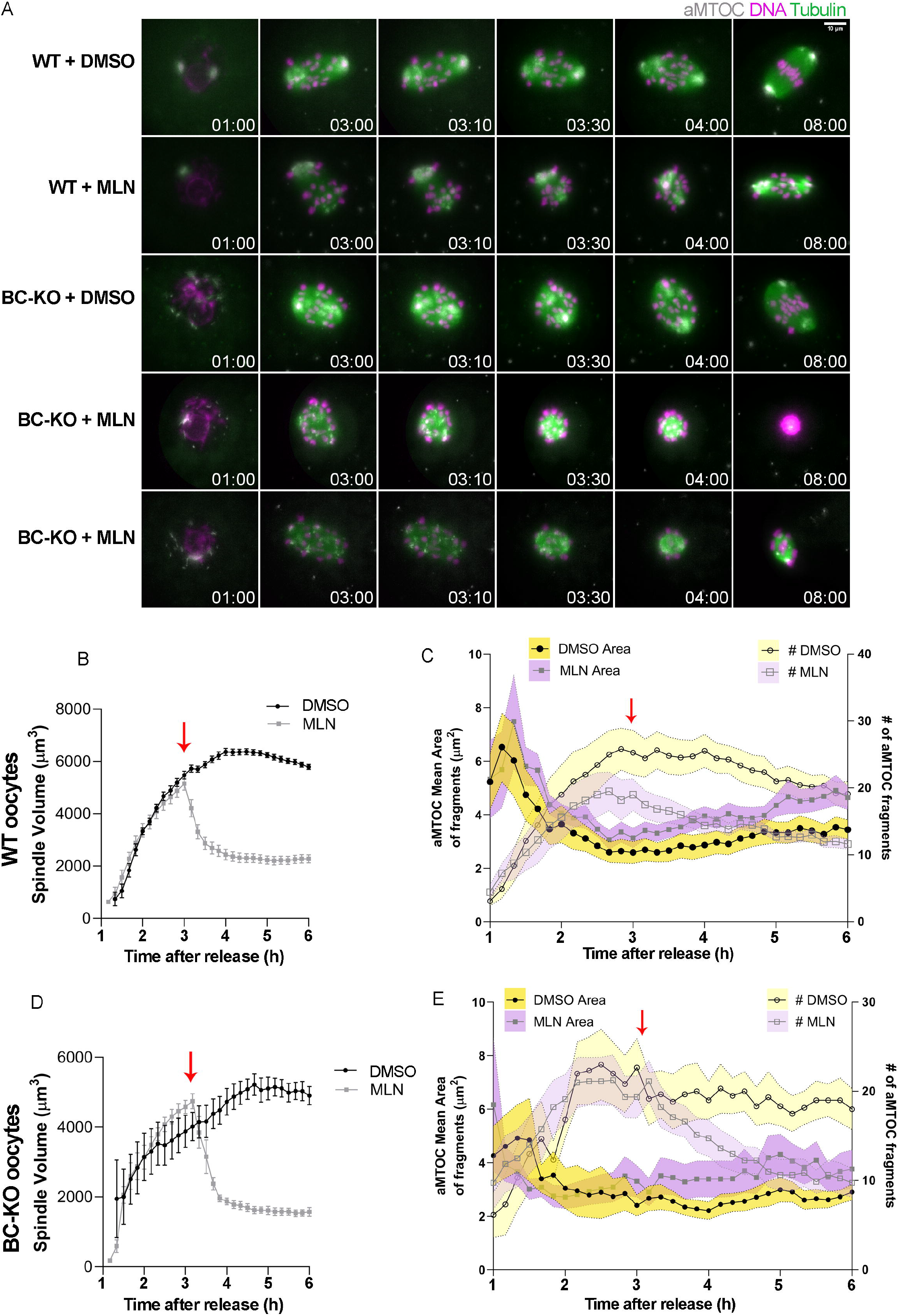
AURKA and AURKC specific functions at late pro-metaphase I in mouse oocytes. (A) Live cell light-sheet imaging of WT and BC-MLN-treated KO oocytes during Early pro-metaphase I; aMTOC (gray), SiR-tubulin (green), DNA (magenta). Time is indicated in h:min; Scale bar 10 μm (B, D) Quantification of spindle volume over time in WT (B) and BC-KO (D), respectively. (C, E) Quantification of area and aMTOC numbers over time in WT (C) and BC-KO (E), respectively. Yellow shadow: DMSO-treated; Purple shadow: MLN-treated. The red arrows indicate MLN addition time. Number of oocytes: WT DMSO: 11; WT MLN: 16; BC-KO DMSO:6; BC-KO MLN:8.

### Polar and chromosomal-targeted AURKA partially rescue spindle defects

Because AURKA localizes to aMTOCs and chromosomes, we asked which localized population is required for spindle building. To answer this question, we generated a mouse strain where oocytes lacked all three AURKs (ABC-KO). This genetic background provides an empty AURK canvas to overexpress different subcellular targeted AURKA fusions without competition or compensation from the others. To confirm that these oocytes lacked all three AURKs, we measured localized AURK activity. In WT oocytes, pABC localized predominantly at spindle poles (active AURKA), and chromosomes (active AURKB/C). No pABC signal was detected in ABC-KO oocytes, consistent with deletion of the three AURKs (Figure S3A, B). We used this model to investigate whether over-expression of WT-AURKA (aMTOC and chromosomes) CDK5RAP2-AURKA (MTOC-AURKA); or H2B-AURKA (Chromatin-AURKA), could rescue the spindle defects in ABC-KO at metaphase I (Figure 4A). First, we tested whether these targeted fusions were active by overexpressing them in ABC-KO oocytes. We observed phosphorylation of INCENP at chromosomes (Figure S4A,B) and CDC25B at aMTOCs (Figure S4C-,D) indicating that these fusion proteins are active at their targeted subcellular localization.

**Figure 4.**
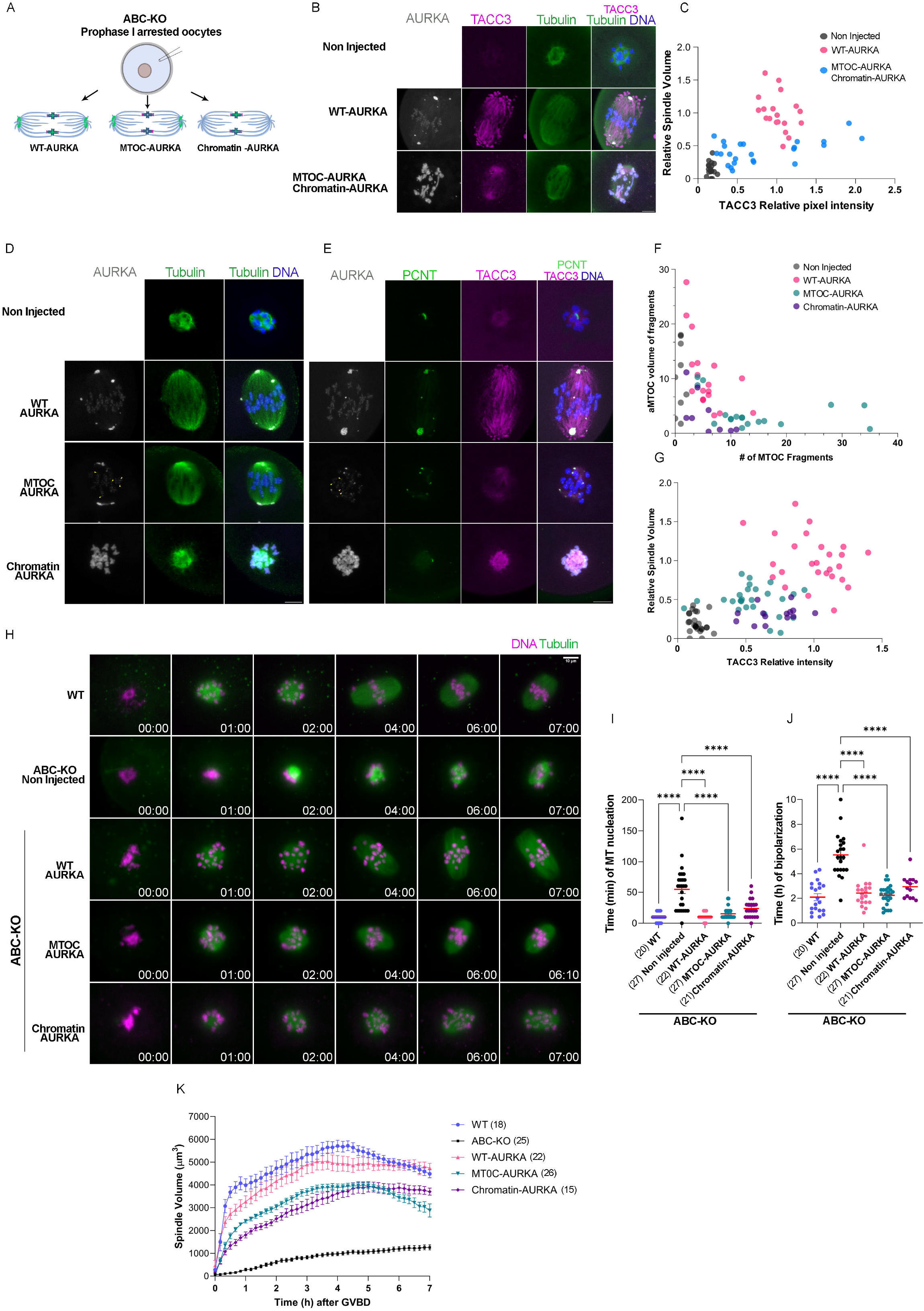
MTOC-AURKA partially rescued spindle phenotypes in ABC-KO oocytes. (A) Schematic of the experimental design. ABC-KO oocytes microinjected with different AURKA fusions. Localization of AURKA is green. (B) Representative confocal images of Metaphase I ABC-KO oocytes expressing the indicated fusions. Non-injected ABC-KO oocytes were control. Oocytes were immuno-stained with TACC3 (magenta), tubulin (green), DAPI (blue). Localization of the targeted AURKA fusion is gray. (C) Quantification of spindle and LISD parameters in (B). Gray dots: Non-injected oocytes; pink dots: WT-AURKA; blue dots: MTOC-AURKA + chromatin-AURKA. (D, E) Representative confocal images of Metaphase I ABC-KO oocytes expressing the indicated fusions immuno-stained with tubulin (green), DAPI (DNA) in (D) and PCNT (green), TACC3 (magenta), DAPI (DNA) in (E). Localization of the targeted AURKA fusion is gray. Yellow arrow indicates MTOC-AURKA at kinetochores. Scale bar: 5µm. (F) Quantification of aMTOC parameters. (G) Quantification of spindle and LISD parameters. Gray dots: Non-injected oocytes; pink dots: WT-AURKA; green dots: MTOC-AURKA, and purple dots: chromatin-AURKA. (H) Live cell light-sheet imaging of WT and ABC-KO oocytes. ABC-KO oocytes expresses the indicated fusion, Sir-tubulin (green), DNA (magenta). (G) Time of MT nucleation and (H) time of spindle bipolarization (One-way ANOVA, **** p<0.0001). Data are represented as mean ± SEM. (K) Spindle volume during meiotic maturation. Time stamp (h:min) is relative to GVBD. In brackets are the number of oocytes analyzed in at least 3 independent experiments. See also Figure S3-S6.

We first asked if expression of both targeted proteins could rescue spindle formation. Non-injected ABC-KO oocytes were used as a reference. ABC-KO oocytes had small spindle volumes and disorganized TACC3 (Figures 4B, C; S5A, B). Upon WT-AURKA overexpression, oocytes were fully rescued, forming bipolar spindles and organized TACC3 (Figures 4B, C; S5A, B). However, when MTOC-AURKA and Chromatin-AURKA were co-expressed, we observed a partial rescue phenotype (Figures 4B, C; S5A, B). TACC3 intensity was largely restored in these oocytes, but spindle volumes were still reduced by ∼50% compared to controls (Figures 4B, C; S5A, B).

To determine which AURKA-localized population was responsible for the partial rescue, we analyzed the same parameters when individual AURKA fusion proteins were overexpressed. We observed that oocytes overexpressing MTOC-AURKA or Chromatin-AURKA also showed intermediate phenotypes (Figure 4D-G). When MTOC-AURKA was overexpressed, the aMTOC fragmentation defect was rescued, because aMTOCs were smaller (Figures 4E,F; S5C, D), but they failed to organize in two defined spindle poles. Although bipolar, spindles were ∼50% smaller (Figures 4D,G; S5E) and TACC3 organization was partially restored compared to controls (Figure 4E,G; S5F). When Chromatin-AURKA was expressed, aMTOC fragmentation was also rescued (Figure 4E, F; S5C, D), but aMTOCs did not separate to form two poles and, these oocytes did not form bipolar spindles. Spindles were 60% smaller (Figures 4D, G; S5E) and TACC3 intensity was 60% reduced compared to WT-AURKA injected controls (Figures 4E, G; S5F).

We next assessed the kinetics of meiotic maturation and spindle formation in ABC-KO oocytes using live-cell imaging. Similar to *Aurka* KO oocytes (Blengini et al., 2021), ABC-KO oocytes formed monopolar or short bipolar spindles (Figure 4H; video S2). We observed that microtubule nucleation onset and spindle bipolarization were significantly delayed (Figure 4H-J). Spindle volume was smaller compared to WT (Figure 4H, K). Expression of each AURKA fusion rescued the timing of microtubule nucleation and spindle elongation (Figure 4H-J). However, spindle volume was only partially rescued in MTOC- or chromatin-AURKA expressing oocytes compared to controls (Fig. 4H, K). These results were consistent with fixed oocyte phenotypes. Taken together, these results suggest that MTOC-localized AURKA is needed for aMTOC fragmentation and spindle bipolarization.

Although the aMTOC-AURKA fusion was designed to localize to aMTOCs, we observed some signal at kinetochores (Figure 4D) where AURKC would normally localize. To resolve specific AURKA vs AURKC functions, we genetically added back AURKC (Figure S3C, D) and evaluated spindle phenotypes in AB-KO oocytes expressing the MTOC-AURKA fusion. When MTOC-AURKA was overexpressed in AB-KO oocytes (AURKC only), we did not observe MTOC-AURKA at kinetochores (Figure S6A). We then compared the level of spindle rescue between AB-KO oocytes and ABC-KO oocytes overexpressing either the WT-AURKA or MTOC-AURKA. Interestingly, there were no significant differences in TACC3 intensity between the genotypes, but there were differences between the AURKA fusion proteins (Figure S6A-C). In both genotypes, when oocytes overexpressed MTOC-AURKA, TACC3 intensity was reduced by 40% compared to controls. There were significant differences in spindle volume both between variants and between genotypes (Figure S6A-C). In AB-KO oocytes overexpressing MTOC-AURKA, spindle volume was reduced by 40% compared to controls. In ABC-KO oocytes, spindle volume was reduced by 50% when MTOC-AURKA was expressed. Although spindle volumes differed between genotypes, we observed that neither AB-KO or ABC-KO oocytes expressing MTOC-AURKA had fully rescued spindle volumes and LISD organization. Taken together, these results suggest that AURKC does not regulate LISD formation but does contribute to a more significant rescue of spindle parameters.

Our data show that neither AURKA fusions fully rescue spindle formation. It is possible that the targeting does not allow for translocation or diffusion along the spindle. In somatic cells, a gradient of AURKA exists extending from centrosomes to metaphase plates. This gradient is important for HEC1 phosphorylation at kinetochores [17], which is critical for spindle elongation and spindle pole formation in mouse oocytes [18]. Therefore, whether Chromatin-AURKA can phosphorylate HEC1, or whether there is a third population of AURKA at kinetochores in oocytes requires further studies. An alternative explanation is that AURKA regulates additional processes that indirectly affect spindle formation. For example, in somatic cells, AURKA localizes to mitochondria [19, 20], and regulates mitochondrial dynamics and ATP production. Whether AURKA has the same role in oocytes is unknown. Another potential AURKA role in oocytes is regulating protein homeostasis. Fully grown oocytes are transcriptionally silent and regulated translation is critical [21]. In *Xenopus* oocytes, AURKA phosphorylates CPEB1, an inhibitor of translation. Phosphorylation of CPEB1 promotes its degradation and translation of maternal mRNA [22]. In mouse oocytes lacking *Aurka*, CPEB1 is not phosphorylated, and levels of total translation are reduced compared to WT [23, 24]. Therefore, by targeting AURKA, we may interfere with AURKA localization to mitochondria or translation activation and affect oocyte energy and protein requirements.

In conclusion, we establish the temporal requirements of AURKA during spindle formation, determine that AURKA localized to aMTOCs is the major population required for spindle organization and suggest roles for AURKC in spindle building during late pro-metaphase I in mouse oocytes. This study raises new questions about non-canonical roles of AURKA during meiotic maturation that are important for understanding the mechanism that ensures oocyte quality that should be further examined.

## Acknowledgments

This work was funded by NIH grant R35 GM136340 to KS and Grant Agency of the Czech Republic grant 23-07532S to DD The authors acknowledge members of the Schindler lab and Devanshi Jain for helpful discussions.

## Author contributions

Conceptualization, C.S.B and K.S.; experimental execution: C.S.B and M.V.; methodology, C.S.B., M.V., D.D., J.S.; data analysis, C.S.B, M.V., D.D.; visualization, C.S.B.; writing original draft, C.S.B. and K.S.; review and editing manuscript, C.S.B, M.V., J.S., D.D., and K.S.; funding,

K.S and D.D.

## Star Method

| REAGENT or RESOURCE | SOURCE | IDENTIFIER |
| --- | --- | --- |
| Antibodies |  |  |
| Mouse monoclonal Pericentrin antibody | BD Biosciences | Cat#611814;RRID: AB_399294 |
| Rabbit TACC3 antibody | Novus Biologicals | Cat# NBP2-67671 |
| Sheep polyclonal Alpha Tubulin antibody | Cytoskeleton | Cat#ATN02; RRID:<br>AB_10708807 |
| Rabbit phosphorylated Aurora A (Thr288), Aurora B (Thr232), Aurora C (Thr198) (D13A11) antibody | Cell Signaling Technology | Cat#2914;RRID:<br>AB_2061631 |
| Rabbit phosphorylated CDC25B (Ser323) antibody | Signalway Antibodies | Cat#11949 |
| phosphorylated INCENP | Gift from Dr. Michael Lampson | Not applicable |
| Donkey anti-mouse Alexa fluor 568 Secondary antibody | Life Technologies | Cat#A10037;<br>RRID:<br>AB_11180865 |
| Donkey anti-rabbit Alexa fluor 647 Secondary antibody | Life Technologies | Cat#A31573;<br>RRID:<br>AB_2536183 |
| Anti-sheep-Alexa-488 Secondary antibody | Life Technologies | Cat#A11015;<br>RRID:<br>AB_2534082 |
| Human polyclonal anti-centromere protein antibody | Antibodies Incorporated | Cat# 15-235;<br>RRID:<br>AB_2939059 |
| Chemicals, peptides, and recombinant proteins |  |  |
| Pregnant Mare Serum Gonadotropin (PMSG) | Lee-Biosolutions<br>Calbiochem | Cat#493-10<br>Cat#367222 |
| Milrinone | Sigma-Aldrich | Cat#M4659 |
| MLN8237 (Alisertib) | Selleck Chemicals | Cat#S1133 |
| MG132 | Selleck Chemicals | Cat#S2619 |
| Dimethyl sulfoxide (DMSO) | Sigma-Aldrich | Cat#472301 |
| Bovine Serum Albumin (BSA) | Sigma-Aldrich | Cat#A4503;<br>Cat#A3311 |
| SiR-tubulin | Spirochrome | Cat#SC002 |
| SiR-DNA | Spirochrome | Cat#SC007 |
| DAPI | Life Technologies | Cat#D1306 |
| Vectashield | Vector Laboratories | Cat#H-1000 |
| L-glutamine | GIBCO | Cat#25030-08 |
| Paraformaldehyde | Sigma-Aldrich | Cat#P6148 |
| CZB culture Media |  |  |
| MEM culture Media | Sigma-Aldrich | Cat#M4655 |
| M2 media | Sigma-Aldrich | Cat#M7167 |
| penicillin–streptomycin | Merck | Cat#P7794,<br>#S1277 |
| Mineral oil | Sigma-Aldrich | Cat#M5310 |
| Sodium pyruvate | Sigma-Aldrich | Cat#P4562 |
| Critical commercial assays |  |  |
| mMESSAGE machine T3 | Ambion, Thermo<br>Fisher Scientific | Cat#AM1348 |
| Poly(A) Tailing Kit | Ambion, Thermo<br>Fisher Scientific | Cat#AM1350 |
| RNeasy Mini Kit | QIAGEN | Cat#74104 |
| Experimental models: Organisms/strains |  |  |
| Mouse: A-KO | Blengini et al [1] |  |
| Mouse: BC-KO | Nguyen et al [2] |  |
| Mouse: AB-KO | This paper |  |
| Mouse: ABC-KO | This paper |  |
| Recombinant DNA |  |  |
| *Aurka-Gfp* | Nguyen et al [2] |  |
| *Aurka-Eyfp* | Blengini et al [1] |  |
| CDK5RAP fr-*Aurka-Eyfp* | This paper |  |
| H2B-*Aurka-Eyfp* | This paper |  |
| *Egfp-Cdk5rap2* | Balboula et al [16] |  |
| *H2b-mCherry* | Kitajima et al [25] |  |
| Software and algorithms |  |  |
| Fiji | National Institute of<br>Health | https://fiji.sc/ |
| Imaris | Oxford instruments | https://imaris.oxinst.com/ |
| Convolutional Neural Network based on the V-net architecture | Milletari et al |  |
| Prim 10 | Graphpad | https://www.graphpad.com/ |

## Resource availability

### Lead contact

Information and requests for resources and reagents should be directed to and will be fulfilled by the lead contact, Karen Schindler.

### Materials availability

Plasmids generated in this study are available without restriction from the lead contact, Karen Schindler. Aurka and Aurkb conditional knockout mice are available without restriction from the lead contact. Aurkc knockout mice were purchased from Taconic and under an MTA license.

### Mouse strains

The different strains of mice used in this work were generated by mating of floxed Aurka (Aurka^f/f^) mice or *Aurkb*^fl/fl^ Gdf9-Cre mice with double knowout mice for AURKB and AURKC *(Aurkb*^f/f^, *Aurkc*^-/-^, Gdf9-Cre mice (BC-KO), described before [1, 2, 26]. After several rounds of mating, we isolated animals that were Aurka^f/f^, Aurkb^f/f^, and Gdf9-Cre mice (AB-KO), and Gdf9-Cre mice (ABC-KO). Moreover, we also used animals Aurkb^f/f^, Aurkc^-/-^, and Gdf9-Cre mice (BC-KO). Control animals (Wild-type (WT)) were Aurka^f/f^, and Aurkb^f/f^ but lacking the Cre recombinase transgene. Mice were housed on a 12–12 h light-dark cycle, with constant temperature and with food and water were provided ad libitum. Animals were maintained in accordance with guidelines of the Institutional Animal Use and Care Committee of Rutgers University (protocol 201702497), the guidelines of National Institutes of Health guidelines, and the policies of the Expert Committee for the Approval of Projects of Experiments on Animals of the Academy of Sciences of the Czech Republic (protocol 106/2020). These regulatory bodies approved all experimental procedures involving the animals. All oocyte experiments were conducted using healthy female mice aged 6–16 weeks. Genotyping was performed before weaning and repeated when the animals were used for experiments as previously described [1, 2].

### Genotyping

Genotyping was performed before weaning and repeated upon use of the animals for experiments for replication and confirmation. LoxP and Cre genotyping was performed by PCR amplification. Primers for *Aurka LoxP* (Forward: 5’-CTGGATCACAGGTGTGGAGT-3’, Reverse: 5’-GGCTACATGCAGGCAAACA-3’), Aurkb LoxP (Forward: 5’-AGGGCCTAATTGCCTCTTGT-3’, Reverse: 5’-GGGCATGAATTCTTGAGTCG-3’), and *Gdf9-Cre* (Forward: 5’-GGCATGCTTGAGGTCTGATTAC-3’, Reverse: 5’-CAGGTTTTGGTGCACAGTCA-3’, Internal control Forward: 5’-CTAGGCCACAGAATTGAAAGATCT-3’, Internal control Reverse: 5’-GTAGGTGGAAATTCTAGCATCATCC-3’). Primers were used at a final concentration of 1 µM, and Taq 2xMaster Mix (NEB, #M0270L) was used following the manufacturer’s protocol. The sizes of products were for *Aurka*: 243 bp for WT and 372 bp for lox/lox transgene; for *Aurkb*: 350 bp for WT and 500 bp for lox/lox transgene, for Cre: 324 bp for no Cre and 200 bp + 324 bp for Cre presence. Deletion of AURKC was detected by a TaqMan copy number assay using primers/probes to detect Neo (Assay #Mr00299300_cn) and Tfrc (for normalization; Assay #4458366). The delta-delta Ct method was used to determine expression levels. PCR conditions are available on request.

### Oocyte collection, maturation, and microinjection

Prophase I-arrested oocytes were collected from ovaries of of 6-16 weeks old females injected 48 hours earlier with 5 I.U. of pregnant mare’s serum gonadotropin (PMSG) (Lee Biosolutions #493–10). Oocytes were collected as described previously [27]. Briefly, ovaries were placed in minimal essential medium (MEM) containing 2.5 μM milrinone (Sigma-Aldrich #M4659) to prevent meiotic resumption and oocytes were isolated by piercing the ovaries with needles. To induce meiotic resumption, oocytes were cultured in milrinone-free Chatot, Ziomek, and Bavister (CZB) medium in an atmosphere of 5% CO2 in air at 37°C. Oocytes were matured for different periods depending on the experimental conditions: to reach early prometaphase I, 3 hours after milrinone wash; for late prometaphase I, 5 hours after milrinone wash; for metaphase I, 7-7.5 hours after milrinone wash, and to evaluate spindle stability we matured the oocytes for 10 hours after milrinone wash.

For microinjection, prophase-arrested oocytes were maintained in CZB supplemented with 2.5 μM milrinone to keep them arrested. Oocytes were microinjected with 100 ng/μl *Aurka-Gfp (WT AURKA),* 100 ng/μl H2B-*Aurka-Eyfp (Chromatin-AURKA),* 100 ng/μl CDK5FRAP fr-*Aurka-Eyfp (MTOC-AURKA), or with* 100 ng/μl H2B-*Aurka-Eyfp and* 100 ng/μl CDK5FRAP fr-*Aurka-Eyfp together. Microinjected oocytes were cultured overnight in* CZB supplemented with 2.5 μM milrinone to allow protein expression prior to the procedures. Subsequently, the oocytes were allowed to mature for 7-7.5 hours to reach metaphase I.

For light-sheet live cell imaging, oocytes were microinjected in M2 medium with ∼10 pl of 50 ng/μl *H2b-mCherry*, 10 ng/μl of *Aurka-Eyfp*, *Aurka*, *H2b-Aurka-Eyfp* and *Aurka-Cdk5rap2-Eyfp*, 100-150 ng/μl *Egfp-Cdk5rap2*, and 75 nM SiR-DNA (Spirochrome, #SC007) or SiR-tubulin (Spirochrome, #SC002) was used for staining chromosomes and spindle, respectively. For protein expression, oocytes were incubated in culture medium supplemented with milrinone for 3 hours after microinjection.

### Immunocytochemistry

After maturation oocytes were immunostained according to a previously described protocol [27]. Briefly, oocytes were fixed in phosphate buffer saline (PBS) containing paraformaldehyde (PFA) 2% for 20 minutes at room temperature. Oocytes were then incubated in permeabilization solution (PBS containing 0.1% (vol/vol) Triton X-100 and 0.3% (wt/vol) BSA) for 20 minutes, followed by 10 minutes in blocking buffer (0.3% BSA containing 0.01% Tween in PBS). Immunostaining was performed by incubating cells in primary antibody overnight in a dark, humidified chamber at 4°C followed by 3 consecutive 10-minute incubations in a blocking buffer. After washing, secondary antibodies were diluted 1:200 in blocking solution and the sample was incubated for 1 hour at room temperature. After washing, the cells were mounted in 10 μL VectaShield (Vector Laboratories, #H-1000) with 4′, 6-Diamidino-2-Phenylindole, Dihydrochloride (DAPI; Life Technologies #D1306; 1:170).

### Live imaging

Oocytes were collected and microinjected in M2 medium (Sigma-Aldrich, #M7167-50ML) containing 2.5 μM milrinone (Sigma-Aldrich) to prevent meiotic maturation. Oocytes were cultured in minimum essential medium (MEM, Merck, #M4655) containing 1.14 mM sodium pyruvate (Merck, #P4562), 4 mg/ml bovine serum albumin (BSA, Merck, #A3311), penicillin– streptomycin (75 U/ml-60 mg/ml, Merck, #P7794, #S1277). Culture and live imaging were performed at 37°C in a 5% CO^2^ atmosphere.

### Plasmid information

Preparation of pYX-EYFP plasmid was described previously (Blengini, 2021). *AURKA* coding sequence was cloned by PCR into pYX-EYFP to create pYX-AURKA-EYFP plasmid. H2B was cloned using NheI restriction sites into pYX-AURKA-EYFP to create pYX-H2B-AURKA-EYFP plasmid. mCDK5RAP2-MBD (amino acids His 1655 – Ser 1822, (NM_145990.4) was cloned into pYX-AURKA-EYFP plasmid by PCR to create pYX-AURKA-EYFP-CDK5RAP2-MBD (original plasmid pGEMHE-mEGFP-mCDK5RAP2 was a gift from Dr. Tomoya Kitajima [25]. CDK5RAP2-MBD fragment corresponds to the microtubule-binding domain (MBD) from human CDK5RAP2 [28].

For *in vitro* transcription, we used a mMESSAGE machine T3 (Ambion, Thermo Fisher Scientific, #AM1348). For polyadenylation of all mRNAs except H2B-mCherry, we used Poly(A) Tailing Kit (Ambion, Thermo Fisher Scientific, #AM1350). All mRNAs were purified using RNeasy Mini Kit (QIAGEN, #74104) and stored at -80°C.

### Drugs and antibodies

Based on a previous study where different Aurora kinase inhibitors were tested in mouse oocytes, to inhibit AURKA, we chose MLN8237 [15] (Alisertib, Selleckchem #S1133). MLN was added to CZB culture media at a final concentration of 1 μM. To avoid the entrance to anaphase I, we inhibited the proteasome by adding 5µM MG132 (Selleck Chemicals, #S2619) to the culture media.

Primary antibodies and concentrations were used as follows: Pericentrin (Pcnt) (mouse, 1:100; BD Biosciences, #611814); TACC3 (Rabbit, 1:100; Novus Biologicals # NBP2-67671), Alpha Tubulin polyclonal (Sheep, 1:100; Cytoskeleton #ATN02); phosphorylated ABC (rabbit, 1:100; Cell Signaling Technology, #2914), phosphorylated CDC25B (1:100; Signalway Antibodies #11949); phosphorylated INCENP (Gift from Michael Lampson), Human anti-ACA (1:30, Antibodies Incorporated #15-234). Secondary antibodies were used at 1:200 for IF experiments: anti-mouse-Alexa-568 (Life Technologies #A10037), anti-rabbit-Alexa-647 (Life Technologies#A31573) and Anti-sheep-Alexa-488 (Life Technologies #A11015)

### Microscopy

For fixed cells, images were captured using a Leica SP8 confocal microscope equipped with 40X. 1.30 NA oil immersion objective. Optical Z-stacks of 1.0 µm step with a zoom of 4.0. In those experiments where pixel intensity was compared, the laser power was kept constant among genotypes or treatments.

For live-cell imaging, oocytes were scanned using the Viventis LS1 Live light-sheet microscope system (Viventis Microscopy Sarl, Switzerland) with a Nikon 25X NA 1.1 detection objective with 1.5 x zoom, as described previously [29]. Thirty-one 2-μm optical sections were taken with a 750x750-pixel image resolution using 10-minute intervals. mClover/EGFP, EYFP, mCherry, and SiR fluorescence were excited by 488, 515, 561, and 638 nm laser lines. To detect mClover/EGFP and EYFP emissions, band-pass filters 525/50 and 539/30 were used. mCherry and SiR emissions were detected using a triple band-pass filter 488/561/640.

### Image analysis

For analysis of TACC3 pixel intensity in fixed oocytes Fiji software [30] was used. We did a maximal projection of the z-stack and we defined the region of interest around the spindle using a free-handed drawing tool, finally, we measured TACC3 intensity in that region of interest using the measurement tool. The volume of spindle and aMTOC fragments in fixed images was performed using Imaris software (Bitplane).

For the videos, image analysis was performed using Fiji software [30]. For basic analysis of meiotic maturation, missegregation, misalignment, MTOC, and spindle bi-polarization max projection from each oocyte was created. For spindle segmentation, we have used a Convolutional Neural Network based on the V-net architecture [31]. The training set consisted of data from 6 experimental datasets, containing between 6 and 20 positions. The annotations were obtained with Fiji, using Gaussian smoothing, followed by constant threshold segmentation and a manual cleanup of the annotation.

### Statistical analysis

All experiments were conducted 3 times, any exception would be clarified in the figure legend. Student T-test, ANOVA one-way or two-way analysis were used to evaluate significant differences between/among groups, using Prism software (GraphPad software). Data is shown as the mean ± standard error of the mean (SEM).

## Supplemental information

### Video S1 – Live cell light-sheet imaging of WT and BC-KO oocytes treated with MLN at early-prometaphase I

MLN was added 3 hours after milrinone wash. aMTOC (gray), DNA (magenta), SiR-tubulin (green). Maximum intensity z-projections are shown. Scale bars: 10 μm. Time points are relative to time after milrinone wash (h:min).

### Video S2 – Live cell light-sheet imaging of WT and ABC-KO oocytes expressing AURKA-targeted fusions

DNA (magenta), SiR-tubulin (green). Maximum intensity z-projections are shown. Scale bars: 10 μm. Time points are relative to time after germinal vesicle breakdown (h:min).

## References

1. Blengini, C.S., Ibrahimian, P., Vaskovicova, M., Drutovic, D., Solc, P., and Schindler, K. (2021). Aurora kinase A is essential for meiosis in mouse oocytes. PLOS Genetics 17, e1009327.

2. Nguyen, A.L., Drutovic, D., Vazquez, B.N., El Yakoubi, W., Gentilello, A.S., Malumbres, M., Solc, P., and Schindler, K. (2018). Genetic Interactions between the Aurora Kinases Reveal New Requirements for AURKB and AURKC during Oocyte Meiosis. Current Biology 28, 3458-+.

3. So, C., Menelaou, K., Uraji, J., Harasimov, K., Steyer, A.M., Seres, K.B., Bucevičius, J., Lukinavičius, G., Möbius, W., Sibold, C., et al. (2022). Mechanism of spindle pole organization and instability in human oocytes. Science 375, eabj3944.

4. Holubcová, Z., Blayney, M., Elder, K., and Schuh, M. (2015). Human oocytes. Error-prone chromosome-mediated spindle assembly favors chromosome segregation defects in human oocytes. Science (New York, N.Y.) 348, 1143–1147.

5. Schuh, M., and Ellenberg, J. (2007). Self-organization of MTOCs replaces centrosome function during acentrosomal spindle assembly in live mouse oocytes. Cell 130, 484–498.

6. Dumont, J., Petri, S., Pellegrin, F., Terret, M.E., Bohnsack, M.T., Rassinier, P., Georget, V., Kalab, P., Gruss, O.J., and Verlhac, M.H. (2007). A centriole- and RanGTP-independent spindle assembly pathway in meiosis I of vertebrate oocytes. J Cell Biol 176, 295–305.

7. Drutovic, D., Duan, X., Li, R., Kalab, P., and Solc, P. (2020). RanGTP and importin beta regulate meiosis I spindle assembly and function in mouse oocytes. EMBO J 39, e101689.

8. Combelles, C.M., and Albertini, D.F. (2001). Microtubule paherning during meiotic maturation in mouse oocytes is determined by cell cycle-specific sorting and redistribution of gamma-tubulin. Dev Biol 239, 281–294.

9. Ma, W., and Viveiros, M.M. (2014). Depletion of pericentrin in mouse oocytes disrupts microtubule organizing center function and meiotic spindle organization. Mol Reprod Dev 81, 1019–1029.

10. Brunet, S., Dumont, J., Lee, K.W., Kinoshita, K., Hikal, P., Gruss, O.J., Maro, B., and Verlhac, M.H. (2008). Meiotic regulation of TPX2 protein levels governs cell cycle progression in mouse oocytes. PLoS One 3, e3338.

11. Clij, D., and Schuh, M. (2015). A three-step MTOC fragmentation mechanism facilitates bipolar spindle assembly in mouse oocytes. Nat Commun 6, 7217.

12. Shuda, K., Schindler, K., Ma, J., Schultz, R.M., and Donovan, P.J. (2009). Aurora kinase B modulates chromosome alignment in mouse oocytes. Mol Reprod Dev 76, 1094–1105.

13. Ding, J., Swain, J.E., and Smith, G.D. (2011). Aurora kinase-A regulates microtubule organizing center (MTOC) localization, chromosome dynamics, and histone-H3 phosphorylation in mouse oocytes. Mol Reprod Dev 78, 80–90.

14. Kratka, C., Drutovic, D., Blengini, C.S., and Schindler, K. (2022). Using ZINC08918027 inhibitor to determine Aurora kinase-chromosomal passenger complex isoforms in mouse oocytes. BMC Res Notes 15, 96.

15. Blengini, C.S., Ik Jung, G., Aboelenain, M., and Schindler, K. (2022). A field guide to Aurora kinase inhibitors: an oocyte perspective. Reproduction 164, V5–v7.

16. Balboula, A.Z., Nguyen, A.L., Gentilello, A.S., Quartuccio, S.M., Drutovic, D., Solc, P., and Schindler, K. (2016). Haspin kinase regulates microtubule-organizing center clustering and stability through Aurora kinase C in mouse oocytes. J Cell Sci 129, 3648–3660.

17. Iemura, K., Natsume, T., Maehara, K., Kanemaki, M.T., and Tanaka, K. (2021). Chromosome oscillation promotes Aurora A-dependent Hec1 phosphorylation and mitotic fidelity. J Cell Biol 220.

18. Courtois, A., Yoshida, S., Takenouchi, O., Asai, K., and Kitajima, T.S. (2021). Stable kinetochore– microtubule ahachments restrict MTOC position and spindle elongation in oocytes. EMBO reports 22, e51400.

19. Grant, R., Abdelbaki, A., Bertoldi, A., Gavilan, M.P., Mansfeld, J., Glover, D.M., and Lindon, C. (2018). Constitutive regulation of mitochondrial morphology by Aurora A kinase depends on a predicted cryptic targeting sequence at the N-terminus. Open Biol 8.

20. Bertolin, G., Bulteau, A.L., Alves-Guerra, M.C., Burel, A., Lavault, M.T., Gavard, O., Le Bras, S., Gagné, J.P., Poirier, G.G., Le Borgne, R., et al. (2018). Aurora kinase A localises to mitochondria to control organelle dynamics and energy production. Elife 7.

21. Richter, J.D., and Lasko, P. (2011). Translational control in oocyte development. Cold Spring Harb Perspect Biol 3, a002758.

22. Han, S.J., Martins, J.P.S., Yang, Y., Kang, M.K., Daldello, E.M., and Conti, M. (2017). The Translation of Cyclin B1 and B2 is Differentially Regulated during Mouse Oocyte Reentry into the Meiotic Cell Cycle. Scientific Reports 7, 14077.

23. Aboelenain, M., and Schindler, K. (2021). Aurora kinase B inhibits aurora kinase A to control maternal mRNA translation in mouse oocytes. Development 148.

24. Kunitomi, C., Romero, M., Daldello, E.M., Schindler, K., and Conti, M. (2024). Multiple intersecting pathways are involved in the phosphorylation of CPEB1 to activate translation during mouse oocyte meiosis. bioRxiv.

25. Kitajima, T.S., Ohsugi, M., and Ellenberg, J. (2011). Complete kinetochore tracking reveals error-prone homologous chromosome biorientation in mammalian oocytes. Cell 146, 568–581.

26. Schindler, K., Davydenko, O., Fram, B., Lampson, M.A., and Schultz, R.M. (2012). Maternally recruited Aurora C kinase is more stable than Aurora B to support mouse oocyte maturation and early development. Proc Natl Acad Sci U S A 109, E2215–2222.

27. Blengini, C.S., and Schindler, K. (2018). Immunofluorescence Technique to Detect Subcellular Structures Critical to Oocyte Maturation. Methods Molecular Biology 1818, 67–76.

28. Wang, Z., Wu, T., Shi, L., Zhang, L., Zheng, W., Qu, J.Y., Niu, R., and Qi, R.Z. (2010). Conserved motif of CDK5RAP2 mediates its localization to centrosomes and the Golgi complex. J Biol Chem 285, 22658–22665.

29. Ferencova, I., Vaskovicova, M., Drutovic, D., Knoblochova, L., Macurek, L., Schultz, R.M., and Solc, P. (2022). CDC25B is required for the metaphase I-metaphase II transition in mouse oocytes. J Cell Sci 135.

30. Schindelin, J., Arganda-Carreras, I., Frise, E., Kaynig, V., Longair, M., Pietzsch, T., Preibisch, S., Rueden, C., Saalfeld, S., Schmid, B., et al. (2012). Fiji: an open-source plarorm for biological-image analysis. Nat Methods 9, 676-682.

31. Milletari, F., Navab, N., and Ahmadi, S.A. (2016). V-Net: Fully Convolutional Neural Networks for Volumetric Medical Image Segmentation. In 2016 Fourth International Conference on 3D Vision (3DV). pp. 565-571.

