## Supplemental figures and legends for "Spatio-temporal requirements of Aurora kinase A in mouse oocytes meiotic spindle building"

**Supplemental information**

**Figure S1. Statistical analyses from figure 1.**

Statistical analyses of WT oocytes treated with MLN at early pro-metaphase I (A-D), late pro-metaphase I (E-H) and Metaphase I (I-L). (A, E, I) Quantification of number of aMTOC fragments. (B, F, J) Quantification of aMTOC volume of fragments. (C, G,K) Quantification of relative spindle volume. (D, H, L) Quantification of TACC3 relative pixel intensity. In brackets are the number of oocytes analyzed in at least 3 independent experiments. Data are represented as mean ± SEM. Unpaired Student t-Test analysis, *p value<0.05; ** p value <0.001; **** p value < 0.0001.

**Figure S2. Statistical analysis from figure 2.**

Statistical analysis of BC-KO oocytes treated with MLN at early pro-metaphase I (A-D), late pro-metaphase I (E-H) and Metaphase I (I-L). (A, E, I) Quantification of aMTOC number. (B, F, J) Quantification of aMTOC volume. (C, G,K) Quantification of relative spindle volume. (D,H,L) Quantification of TACC3 relative pixel intensity. In brackets is represented the number of oocytes analyzed in at least 3 independent experiments. Data are represented as mean ± SEM. Unpaired Student t-Test analysis, *p value<0.05; ** p value <0.001; **** p value < 0.0001.

**Figure S3. Genotype confirmation of ABC-KO and AB-KO oocytes.**

Representative confocal images of metaphase I oocytes from WT and ABC-KO (A) and AB-KO (C) females immuno-stained to detect pAURKA/B/C (pABC) (gray), ACA (magenta), tubulin (green), and DAPI (blue). Scale bar: 5µm. (B, D) Quantification of pABC relative pixel intensity of (A) and (C). In brackets is the number of oocytes analyzed from 1 mouse. Unpaired Student t-Test analysis ****p<0.0001. Data are represented as mean ± SEM.

**Figure S4. Targeted AURKA fusion proteins are active at chromosomes and aMTOCs**. (A) Representative confocal images of ABC-KO oocytes expressing either WT-AURKA or chromatin-AURKA immuno-stained with pINCENP (gray), DAPI (blue) and the overexpressed AURKA localization (green). (B) Quantification of pINCENP relative pixel intensity from (A). (C) Representative confocal images of ABC-KO oocytes expressing either WT-AURKA or MTOC-AURKA immuno-stained with pCDC25B (gray), DAPI (blue) and the overexpressed AURKA localization (green). (D) Quantification of pCDC25B relative pixel intensity from (C). In brackets is the number of oocytes analyzed. Unpaired Student t-Test analysis, *** p value <0.001. Data are represented as mean ± SEM.

**Figure S5. Statistical analyses from figure 4.**

(A) TACC3 relative pixel intensity and (B) relative spindle volume in ABC-KO oocytes expressing the indicated fusions. (C) Number of aMTOCs, (D) aMTOC volume, (E) relative spindle volume and, (F) TACC3 relative pixel intensity in ABC-KO oocytes expressing the indicated fusions. In brackets is the number of oocytes analyzed in at least 3 independent experiments. One-way ANOVA *p value<0.05; ** p value <0.01; ***p value< 0.001; **** p value < 0.0001. Data are represented as mean ± SEM.

**Figure S6. AURKC prevents MTOC-AURKA from binding chromosomes.**

(A) Representative confocal images of AB-KO and ABC-KO oocytes expressing either WT-AURKA or MTOC-AURKA immuno-stained with tubulin (green), TACC3 (magenta), DAPI (blue) and AURKA (gray). Scale bar: 5µm. (B) Quantification of relative spindle volume of (A) Two-way ANOVA, genotype factor p=0.0010; targeted AURKA fusion factor p<0.0001; interaction p=0.1471). (C) Two-way ANOVA, genotype factor p=0.0713; targeted AURKA fusion factor

$p<0.0001$; interaction $p=0.1491$). In brackets is the number of oocytes analyzed in at least 3 independent experiments. Data are represented as mean ± SEM.

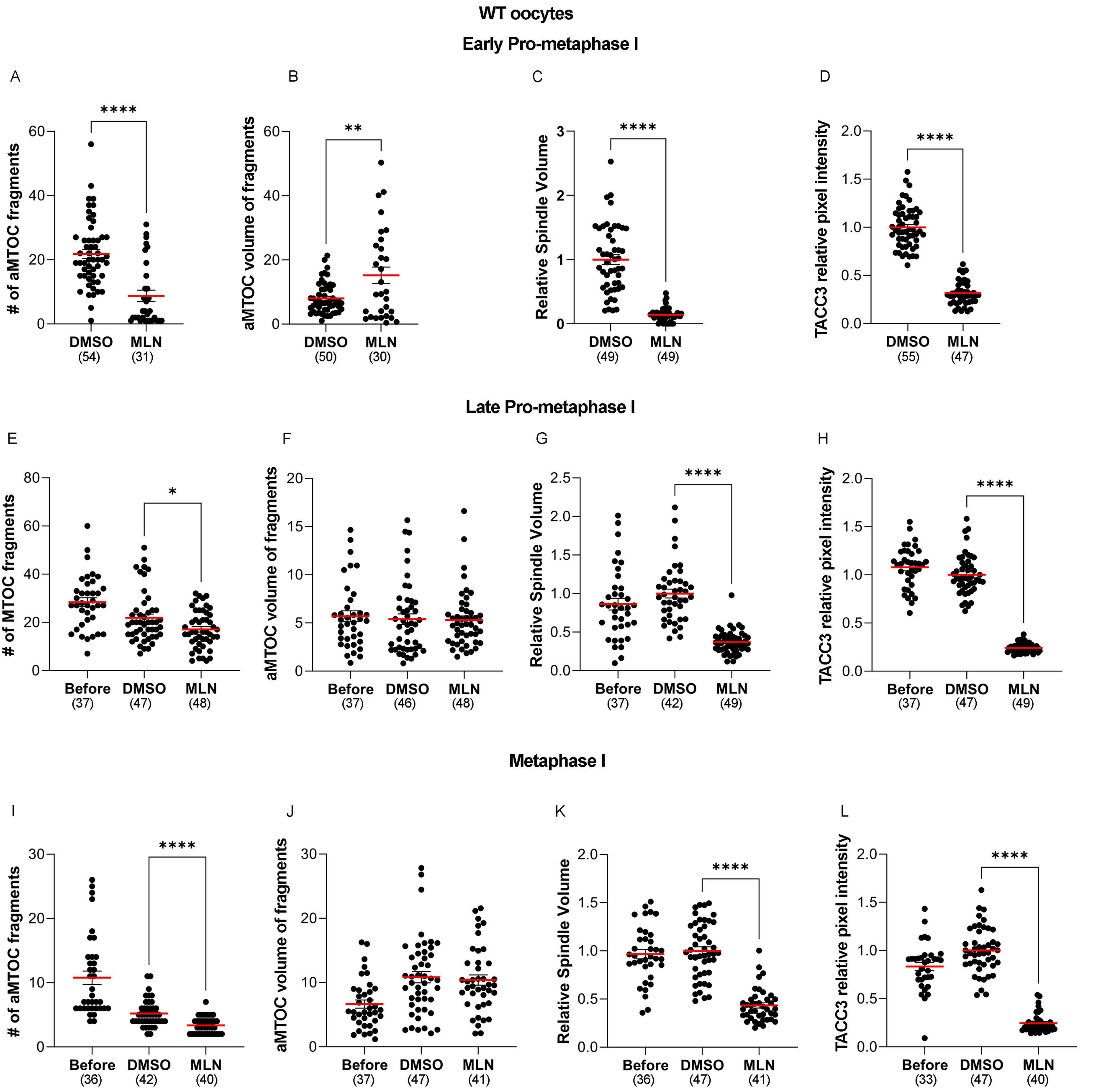
WT oocytes
Early Pro-metaphase I
A
B
C
D
### of aMTOC fragments
aMTOC volume of fragments
Relative Spindle Volume
TACC3 relative pixel intensity
DMSO (54) MLN (31)
DMSO (50) MLN (30)
DMSO (49) MLN (49)
DMSO (55) MLN (47)
Late Pro-metaphase I
E
F
G
H
### of MTOC fragments
aMTOC volume of fragments
Relative Spindle Volume
TACC3 relative pixel intensity
Before (37) DMSO (47) MLN (48)
Before (37) DMSO (46) MLN (48)
Before (37) DMSO (42) MLN (49)
Before (37) DMSO (47) MLN (49)
Metaphase I
I
J
K
L
### of aMTOC fragments
aMTOC volume of fragments
Relative Spindle Volume
TACC3 relative pixel intensity
Before (36) DMSO (42) MLN (40)
Before (37) DMSO (47) MLN (41)
Before (36) DMSO (47) MLN (41)
Before (33) DMSO (47) MLN (40)

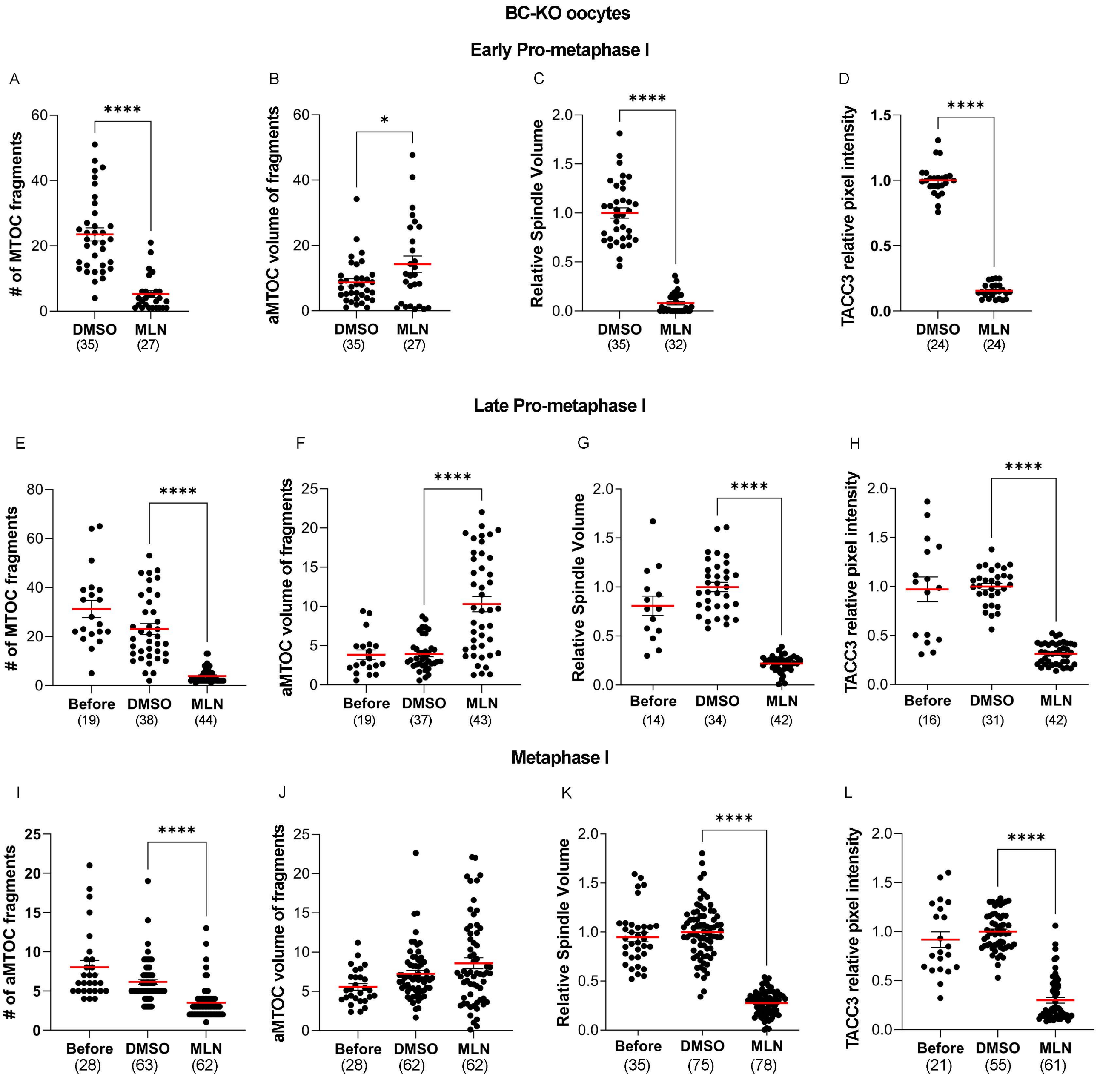
BC-KO oocytes
Early Pro-metaphase I
A
### of MTOC fragments
DMSO (35)
MLN (27)
****
B
aMTOC volume of fragments
DMSO (35)
MLN (27)
*
C
Relative Spindle Volume
DMSO (35)
MLN (32)
****
D
TACC3 relative pixel intensity
DMSO (24)
MLN (24)
****
Late Pro-metaphase I
E
### of MTOC fragments
Before (19)
DMSO (38)
MLN (44)
****
F
aMTOC volume of fragments
Before (19)
DMSO (37)
MLN (43)
****
G
Relative Spindle Volume
Before (14)
DMSO (34)
MLN (42)
****
H
TACC3 relative pixel intensity
Before (16)
DMSO (31)
MLN (42)
****
Metaphase I
I
### of aMTOC fragments
Before (28)
DMSO (63)
MLN (62)
****
J
aMTOC volume of fragments
Before (28)
DMSO (62)
MLN (62)
K
Relative Spindle Volume
Before (35)
DMSO (75)
MLN (78)
****
L
TACC3 relative pixel intensity
Before (21)
DMSO (55)
MLN (61)
****

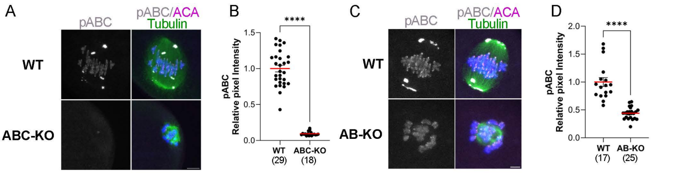
A
pABC
pABC/ACA
Tubulin
WT
ABC-KO
B
****
pABC
Relative pixel Intensity
1.5
1.0
0.5
0.0
WT
(29)
ABC-KO
(18)
C
pABC
pABC/ACA
Tubulin
WT
AB-KO
D
****
pABC
Relative pixel Intensity
2.0
1.5
1.0
0.5
0.0
WT
(17)
AB-KO
(25)

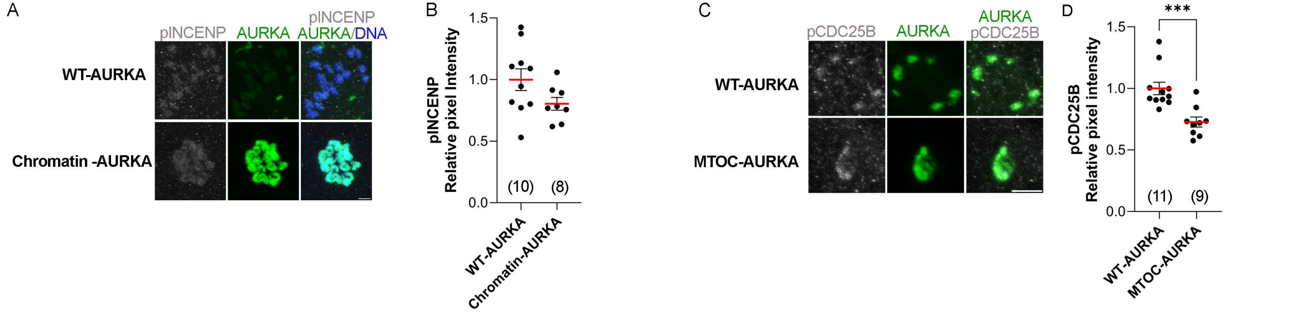

A
pINCENP
AURKA
pINCENP
AURKA/DNA
WT-AURKA
Chromatin -AURKA
B
pINCENP
Relative pixel Intensity
1.5
1.0
0.5
0.0
(10)
(8)
WT-AURKA
Chromatin-AURKA
C
pCDC25B
AURKA
AURKA
pCDC25B
WT-AURKA
MTOC-AURKA
D
***
pCDC25B
Relative pixel intensity
1.5
1.0
0.5
0.0
(11)
(9)
WT-AURKA
MTOC-AURKA

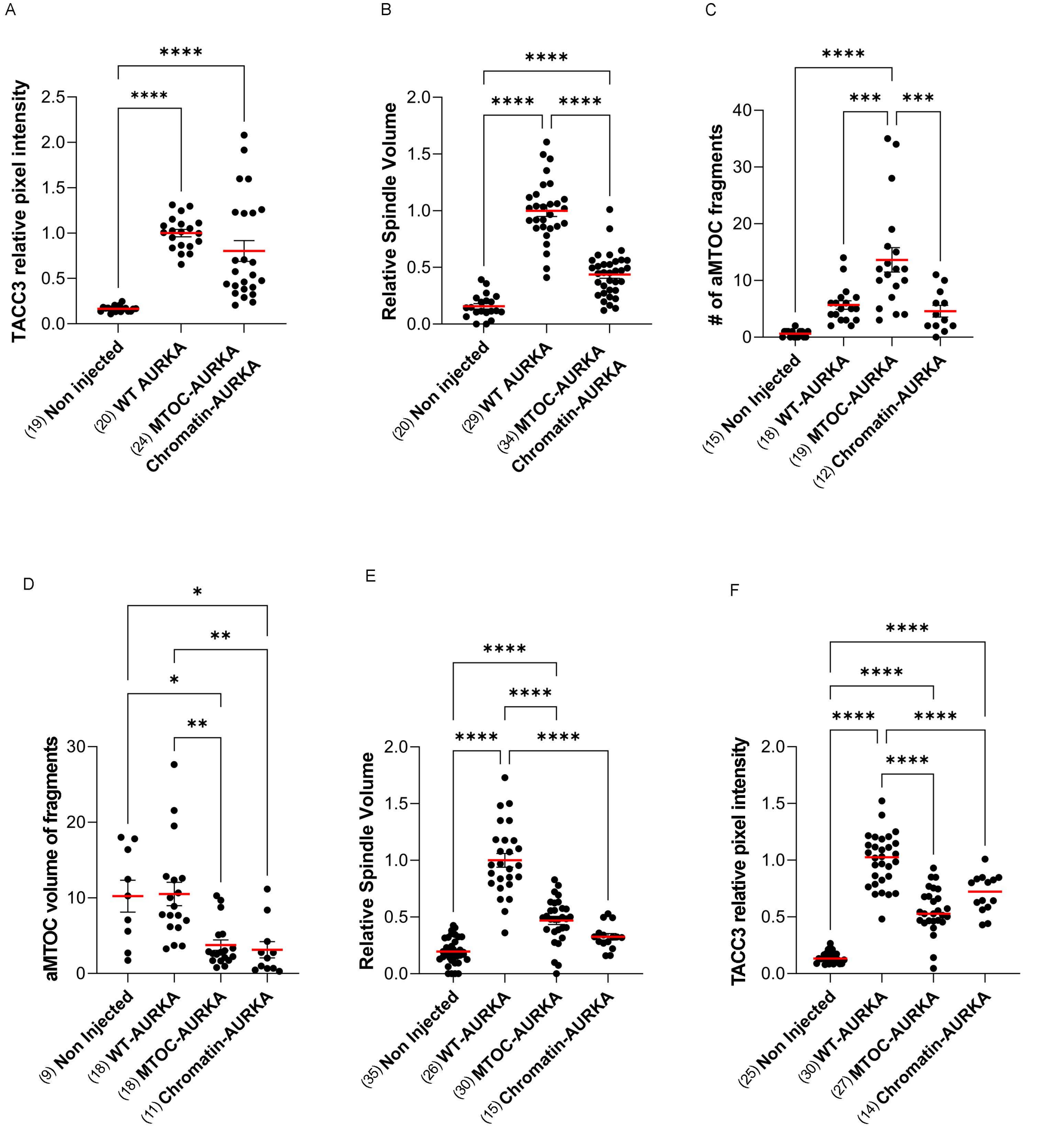
A
TACC3 relative pixel intensity
0.0
0.5
1.0
1.5
2.0
2.5
****
****
(19) Non injected
(20) WT AURKA
(24) MTOC-AURKA
Chromatin-AURKA
B
Relative Spindle Volume
0.0
0.5
1.0
1.5
2.0
****
****
****
(20) Non injected
(29) WT AURKA
(34) MTOC-AURKA
Chromatin-AURKA
C
### of aMTOC fragments
0
10
20
30
40
****
***
***
(15) Non Injected
(18) WT-AURKA
(19) MTOC-AURKA
(12) Chromatin-AURKA
D
aMTOC volume of fragments
0
10
20
30
*
**
*
**
(9) Non Injected
(18) WT-AURKA
(18) MTOC-AURKA
(11) Chromatin-AURKA
E
Relative Spindle Volume
0.0
0.5
1.0
1.5
2.0
****
****
****
****
(35) Non Injected
(26) WT-AURKA
(30) MTOC-AURKA
(15) Chromatin-AURKA
F
TACC3 relative pixel intensity
0.0
0.5
1.0
1.5
2.0
****
****
****
****
****
(25) Non Injected
(30) WT-AURKA
(27) MTOC-AURKA
(14) Chromatin-AURKA

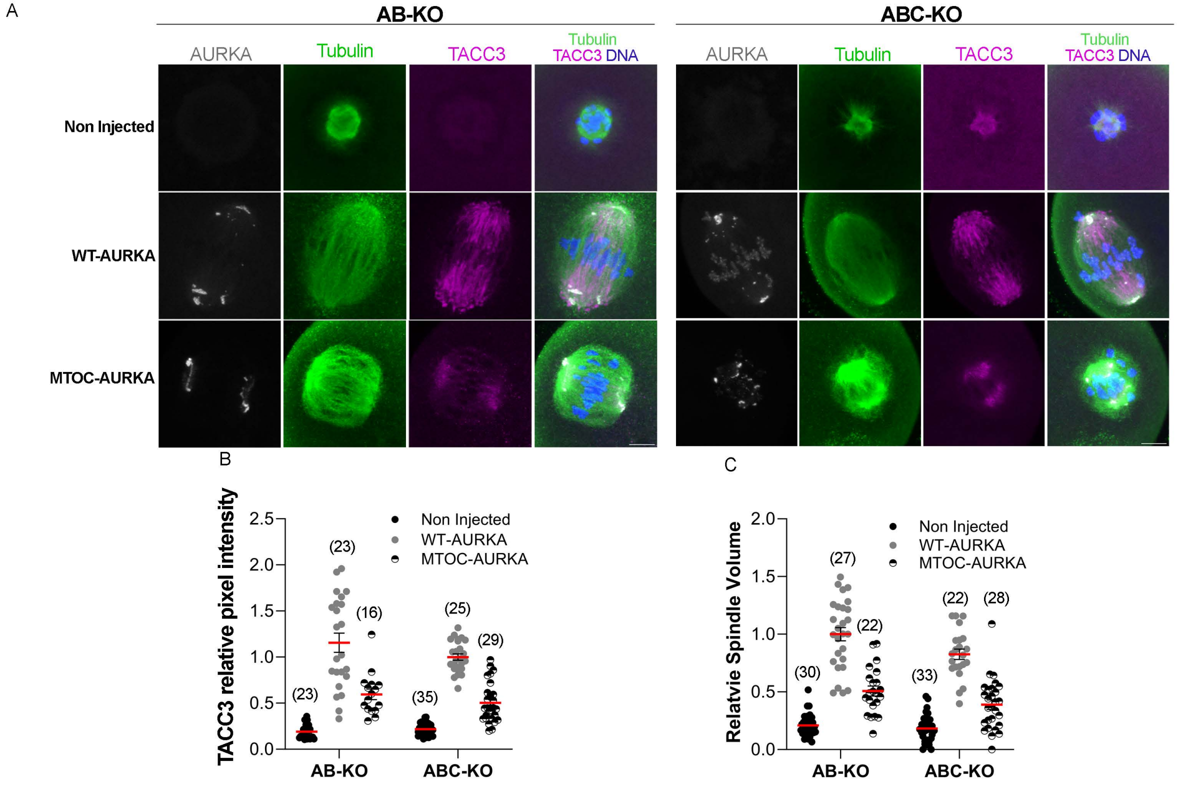
A
AB-KO
ABC-KO
AURKA
Tubulin
TACC3
Tubulin TACC3 DNA
Non Injected
WT-AURKA
MTOC-AURKA
B
TACC3 relative pixel intensity
2.5
2.0
1.5
1.0
0.5
0.0
Non Injected
WT-AURKA
MTOC-AURKA
(23)
(23)
(16)
(35)
(25)
(29)
AB-KO
ABC-KO
C
Relatvie Spindle Volume
2.0
1.5
1.0
0.5
0.0
Non Injected
WT-AURKA
MTOC-AURKA
(30)
(27)
(22)
(33)
(22)
(28)
AB-KO
ABC-KO
